# Robust non-invasive detection of hyperglycemia in mouse models of metabolic dysregulation: the novel Urination Index biomarker

**DOI:** 10.1101/2025.04.02.646666

**Authors:** Sebastian Brachs, Morten Dall, Leonie-Kim Zimbalski, Yohan Santin, Christian Oeing, Knut Mai, Angelo Parini, Stefano Gaburro, Thomas Svava Nielsen

## Abstract

Blood glucose is one of the most essential parameters in metabolic research. Yet, accurate blood glucose monitoring in mouse models of diabetes is challenging due to the significant stress associated with the measurements and the variability in diabetes development among experimental mouse models. This variability necessitates frequent blood glucose measurements, which only provide intermittent data and may not accurately reflect continuous metabolic changes.

To address these issues, we have utilized the Tecniplast DVC system to monitor bedding moisture, enabling the detection of increased urination (polyuria), a primary symptom of diabetes. Polyuria is a hallmark of (undiagnosed/untreated) diabetes, and we revealed high correlations between bedding moisture and blood glucose during hyperglycemia. Thus, our developed algorithm enhances animal welfare by reducing the need for invasive blood glucose tests and enabling non-invasive, continuous assessment of hyperglycemia onset, progression, and severity directly within the mice’s home cage. Continuous monitoring of polyuria allows detailed analysis of temporal and circadian urination patterns and enables assessment of the efficacy of glucose-lowering interventions, which is crucial in developing new pharmacological treatments. We propose that this innovative approach of a novel digital biomarker, the Urination Index (UI), offers a significant advance in the methodology for diabetes research in mouse models, improves animal welfare by reducing the need for invasive blood glucose tests, and enhances the reliability of data and the quality of life for the animals involved.

## Introduction

Diabetes mellitus (DM) is a major global health problem affecting >800 million people and causing severe challenges for healthcare systems^1^. Chronic hyperglycemia (blood glucose >10 mM) is a critical factor^2^, accounting for 43% of all deaths under 70 years^3^. Therefore, blood glucose assessment is indispensable in animal models used in metabolic research and pharmacology, but it is invasive with known drawbacks^4^. In mice, accurate monitoring of blood glucose is a major challenge, primarily due to the sampling stress triggering hyperglycemia, and due to the inherent variability in disease progression across and within models^4–9^. Traditional sampling using glucometers is invasive, labor-intensive and offers only intermittent glucose snapshots that fail to capture dynamic fluctuations^10^. Continuous glucose monitoring devices (CGM) are cost-intensive and requires specialized surgical skills and anesthesia, which has significant glucose regulating effects^11–13^. Thus, reliable glucose monitoring is difficult in mice and the measurements may affect experimental outcomes^14,15^.

Excessive urination, known as polyuria, is a first clinical manifestation and common symptom of hyperglycemia in undiagnosed or poorly managed DM accompanied by polydipsia and polyphagia^16^. Voiding behavior, the pattern and frequency of urination, is regulated by a circadian rhythm and is known to be altered in DM^17–19^. The most common methods to analyze urine and voiding behavior in laboratory animals are void spot assays, catheterization and metabolic cages, which are all associated with stress^20–25^.

As hyperglycemia has a complex diurnal rhythm and is a main driver of polyuria, we hypothesized that monitoring of urination patterns could provide valuable insights about blood glucose levels in mouse models of DM. Therefore, we implemented an innovative approach using bedding moisture as a measure of polyuria for non-invasive continuous monitoring of hyperglycemia. To assess bedding moisture, we utilized the DVC (Digital Ventilated Cage, Tecniplast) sensing technology. The DVC system automatically records home cage data for multiple parameters for daily routine tasks in the animal husbandry management and animal welfare support^26,27^. However, multiple approaches used the DVC to derive further parameters, such as cage aggression, behavioral traits, activity and circadian profiles^28–33^.

In this work, we assessed increased cage urination (polyuria) in mouse models of type-1 (T1DM) and type-2 DM (T2DM) by detection of bedding moisture using the DVC system and were able to correlate it with polydipsia, polyphagia and hyperglycemia. We developed a novel digital biomarker, the Urination Index (UI), for non-invasive assessment of urination and developed an app to analyze it (https://cbmr-rmpp.shinyapps.io/UrinatoR/). This approach enables sample-free continuous tracking of the progression of hyperglycemia within the home cage. Our innovative approach adheres to the 3R principles by reducing invasive sampling, handling and, therefore, stress, improving animal welfare and increasing data reliability. Furthermore, unlike intermittent measurements, longitudinal and circadian urination patterns can be identified in high temporal resolution, including during the dark phase, when mice are active. Finally, we could reverse the polyuria phenotype by pharmacological blood glucose-lowering intervention in hyperglycemic streptozotocin-treated mice. This real-time remote monitoring of disease onset, progression and severity in the home cage environment enables timing of interventions or treatments according to the onset of hyperglycemia, limiting model variability and improving outcomes of metabolic, diabetic and pharmacological research.

## Results

### Basic urination algorithm

The DVC detects increasing bedding moisture as a decline in electromagnetic field strength and records this as the Bedding Status Index (BSI, Fig. 1a). To obtain a metric that increases with added moisture, the BSI was converted into inverse incremental changes for each data point (Fig. 1b). After removal of signal corresponding to bedding changes (Fig. 1c), the Urination Index (UI) was calculated as the cumulative development of inverted and processed BSI (Fig. 1d). Hence, UI is a cumulative metric, the inverse of the BSI with the discontinuities of bedding changes removed, having the same scale and arbitrary unit as the BSI.

**Fig. 1:**
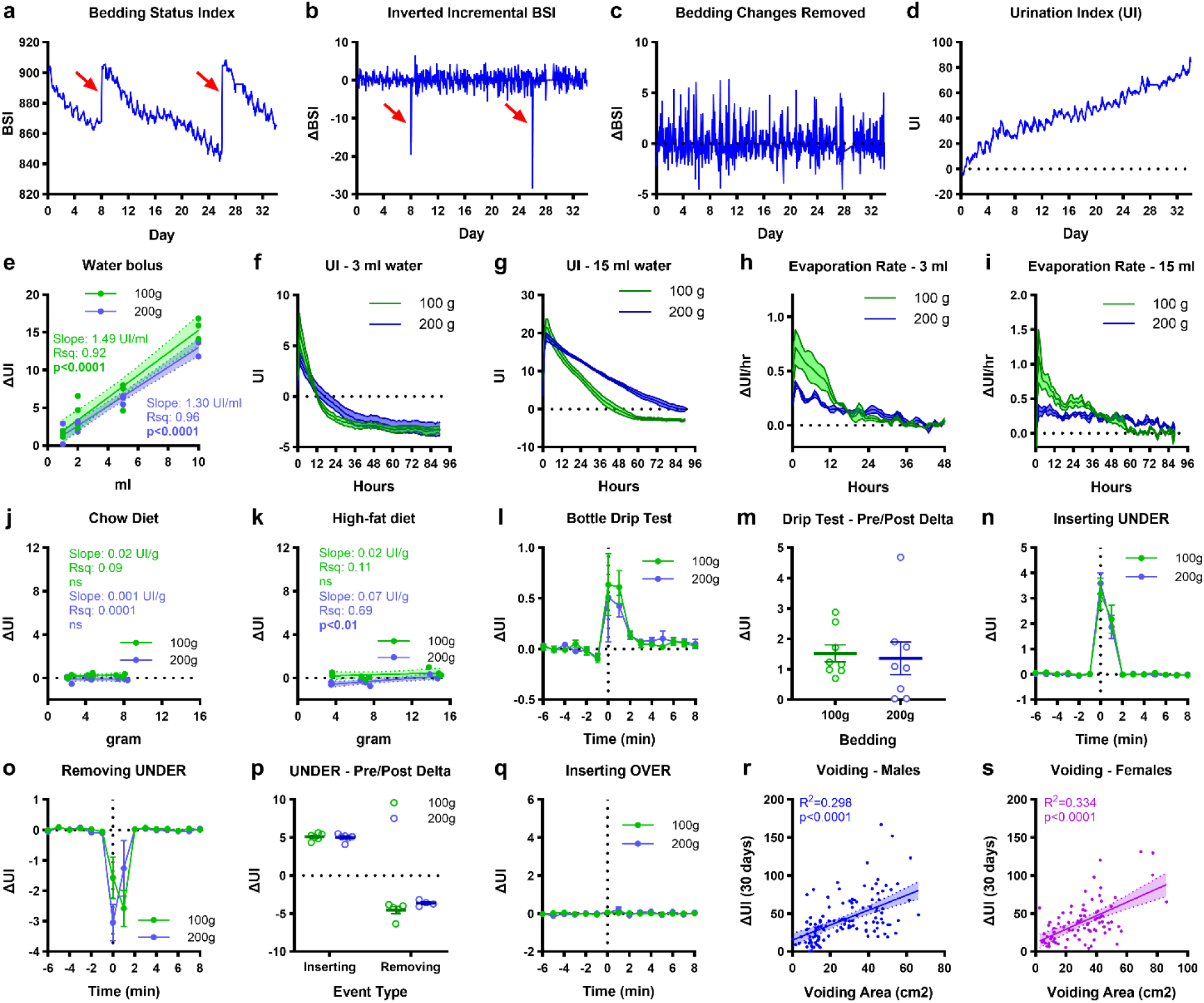
Derivation of Urination Index (UI) algorithm and in vitro validation. **a,** The DVC Bedding Status Index (BSI) from one cage. Two bedding changes are indicated (arrows). **b**, Conversion of the BSI to incremental changes and inversion of the sign. **c**, Incremental changes after removal of bedding changes. **d**, UI generated by accumulation of the processed incremental data. **e**, Dose-response test of injecting water in cages without mice.1,2, 5, and 10ml injected in cages with 100g or 200g of bedding (n=3). 95% CI (shaded area) and regression results are indicated. **f-g**, UI change during evaporation of 3ml (**f**) or 15ml (**g**) of water from bedding (mean±SEM, n=3). **h-i**, Rate of evaporation of 3ml (**h**) and 15ml (**i**) of water, calculated as the slope of **f-g** (mean±SEM, n=3). **j-k**, UI response and regression results to adding 1, 2, and 4 chow diet pellets (n=3) (**j**) or high-fat diet pellets (n=3) (**k**). **l,** UI response to bottle dripping during cage handling (mean±SEM, n=8). **m**, Total change in UI after a cage handling event (mean±SEM, n=8). **n**, UI response to insertion of a cage in the slot under the monitored cage (mean±SEM, n=6). **o**, UI response to removal of a cage from the slot under the monitored cage (mean±SEM, n=6). **p**, Total change in UI when a cage is inserted or removed under the monitored cage (mean±SEM, n=6). **q**, UI response to insertion of a cage in the slot above the monitored cage (mean±SEM, n=6). **r-s**, Correlation and 95% CI (shaded area) between the 30-day UI increase and the total urine spot area during voiding test in male (n=123, **r**) and female (n=87, **s**) Swiss mice from the INSPIRE cohort.

### “In vitro” tests of water, diet, and cage-handling effects

To establish the relationship between UI and added water or diet in cages, we conducted controlled “in vitro” experiments without animals using cages with varying amounts of bedding. The UI signal showed a strong linear relationship with the volume of added water (Fig. 1e). Since slopes were not dependent on bedding amount (p>0.05), we could estimate the average effect of water on the signal to be 1.4UI/ml. Conversely, evaporation rates were dependent on bedding amount. With 3ml and 15ml volume injected, the UI signal had a steeper initial decline and reached a plateau sooner in cages with 100g than with 200g bedding (Fig. 1f-g). Calculating the evaporation rate revealed that irrespective of initial water content, 200g bedding promoted a more constant evaporation rate than 100g, indicating that more bedding stabilized the UI signal (Fig. 1h-i). Unlike water, chow or HFD pellets had minimal effects on the UI signal (0.001-0.07UI/g, i.e., <5% of an equivalent mass of water, Fig. 1j-k), suggesting that diet spillage or shredding is unlikely to contribute to the estimate of urination. Since there is a risk of bottle spillage when handling cages, we simulated a standard cage-handling procedure where cages were removed, opened, closed, and returned to the DVC (Fig. 1l). Indeed, dripping can cause a substantial UI change, with an overall average of 1.4±0.3UI units in both bedding groups (Fig. 1m). However, the inter-cage differences varied considerably between 0.02-4.7UI units (corresponding to 0.01-3.4ml). As the DVC electrodes are omnidirectional, they are also expected to detect the proximity of water in a bottle immediately below. Consequently, we inserted and removed surrounding cages and indeed, inserting a cage with a full bottle below caused a substantial UI increase in the cage above (Fig. 1n). Vice versa, a UI reduction was observed in the upper cage removing that below (Fig. 1o). The insertion/removal effect below was equal and opposite, both causing an offset of 5UI units (Fig. 1p). As expected, inserting/removing a cage above did not affect the UI of the cage below (Fig. 1q). Altogether, our tests indicate that UI is a reliable and selective measure of changes in bedding fluid content, but also that exclusion of measurements associated with cage-handling and insertion/removal events is crucial in data processing.

### UI correlates with voiding behavior

We used data of male and female mice from the INSPIRE cohort^33,34^ to correlate the UI to results of the voiding behavior test. Here, we found a positive correlation between total voiding spot area and UI indicating that the UI is a valid measure of urinary output (males: R²=0.55, p<0.0001, Fig. 1r, females: R²=0.58, p<0.0001, Fig. 1s).

### Housing density scales with increasing density

To evaluate our approach, we analyzed the linear scalability of the UI performing a housing density test with male and female mice (Fig. 2a-b). BSI was converted to UI per cage with cage density of 1, 2, 3, 4, or 5 mice (Fig. 2c-d). Thereafter, we normalized by housing density to derive UI per mouse (Fig. 2e-f). The urination rate per cage calculated as the slope of the UI curve showed a strong linear correlation with cage density in both sexes (males: 0.63UI/mouse/day, R^2^=0.78, p<0.05, Fig. 2g, females: 0.41UI/mouse/day, R^2^=0.96, p<0.01, Fig. 2h). After housing density normalization, this correlation was no longer present for males (Fig. 2g). Although a slight correlation was still present after normalization in females (0.04UI/mouse/day, R^2^=0.84, p<0.05, Fig. 2h), these results demonstrate the linear scalability and robustness of our approach across different housing densities.

**Fig. 2:**
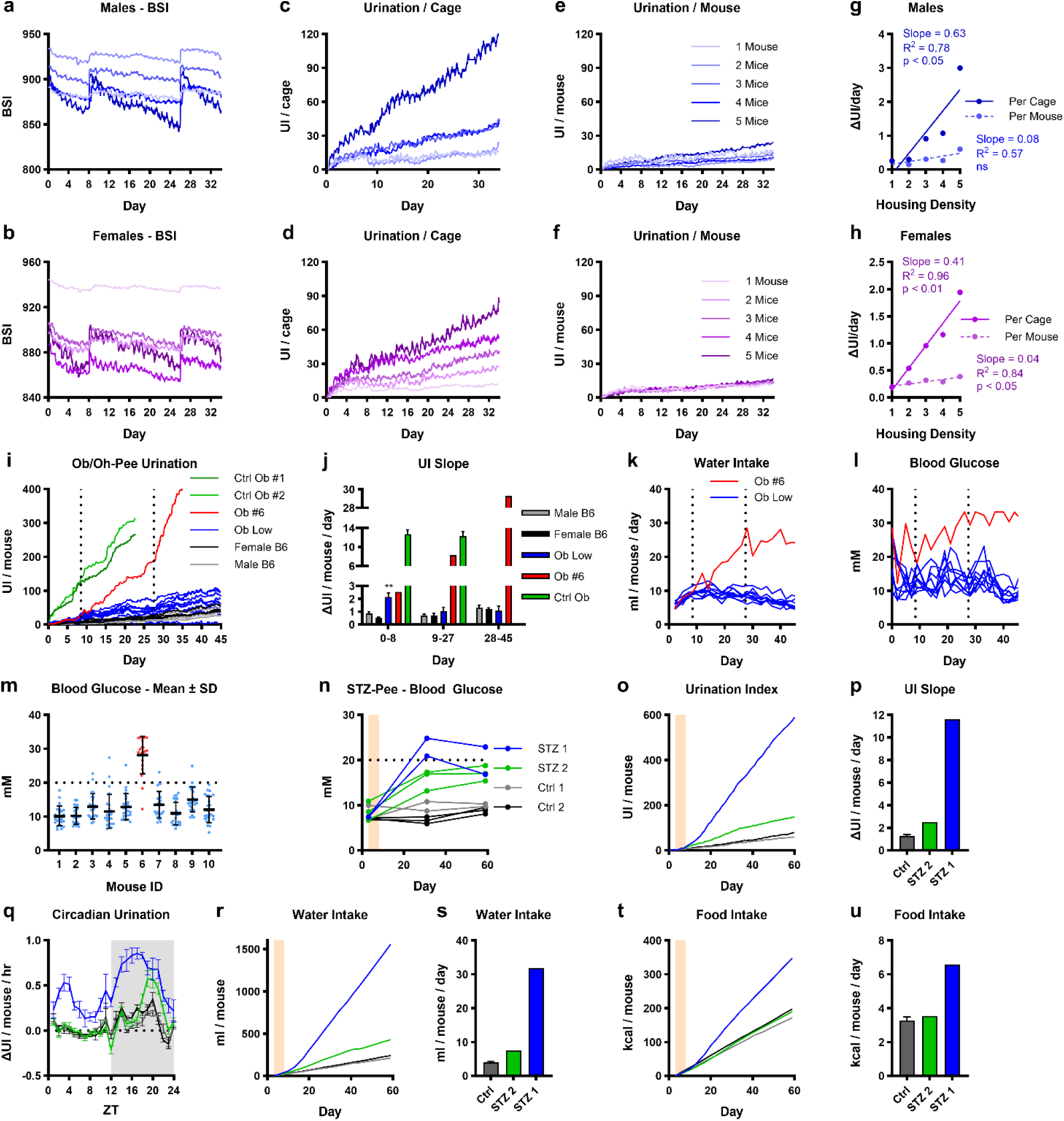
Housing density and diabetes pilot studies. **a-b,** Bedding Status Index (BSI) from 1-5 male (**a**) and 1-5 female (**b**) CD1 mice (n=1). **c-d**, UI for each cage of male (**c**) and female (**d**) mice. **e-f**, UI normalized for housing density of male (**e**) and female (**f**) mice. **g-h**, Relationship between UI and housing density in male (**g**) and female (**h**) mice before and after normalization. **i**, Ob/Oh-Pee study UI normalized for housing density in male (n=4) and female (n=4) C57BL/6, non-responder ob/ob (Ob-Low, n=9), responder ob/ob mouse (Ob-#6, n=1), and positive control ob/ob (Ctrl Ob-#1, Ob-#2 n=2). Dashed lines indicate time of response changes in Ob-#6. **j**, Average daily urination rate in each study period (mean±SEM, **p<0.01 vs. male B6). **k-l**, Time course of individual daily water intake (**k**) and blood glucose (**l**) in Ob Low mice and Ob #6. **m**, Blood glucose for each mouse in Ob Low (blue) and Ob #6 (red) (mean±SD). **n**, Blood glucose time course for each mouse in STZ-Pee pilot study (blue and green: STZ-treated cages, grey and black: control cages). STZ was dosed on day 3-8 (shaded area) **o**, UI normalized for housing density. **p**, Average daily urination rate (UI slope day 10-60, Ctrl: mean±SEM, n=2). **q**, Circadian urination pattern (day 10-60, individual cages). **r**, cumulative water intake. **s**, average daily water intake (water intake slope day 10-60, Ctrl: mean±SEM, n=2). **t**, Cumulative food intake. **u**, average daily food intake (food intake slope day 10-60, Ctrl: mean±SEM, n=2).

### Ob/Oh-Pee Study

As initial proof of concept, we monitored two severely hyperglycemic ob/ob mice (Ctrl Ob-#1, Ob-#2), a T2DM model, with constant blood glucose >33.3mmol/L. These mice showed dramatically increased UI compared to wildtype C57BL/6J (WT) controls (Fig. 2i). Consequently, we conducted a study with 10 individually housed ob/ob males, monitored from 3 weeks of age, to study UI progression during hyperglycemia development. Compared to WT, 9 ob/ob mice had only mildly elevated UI (Fig. 2i). However, Ob-#6 presented with severe polyuria starting from day 9 (Fig. 2i). On day 28, it also developed a stereotypic behavior of excessive grooming on the water nozzle, causing spillage and further exacerbated the increase in UI. By calculating the slope of UI curves, we derived the average daily ΔUI for each phase of the study (Fig. 2j). In the first 8 days, the ΔUI of 9 ‘Ob-Low’ mice was 2.5-fold elevated (Ob-Low: 2.2±0.3UI/mouse/day, male WT: 0.9±0.1UI/mouse/day, p<0.01). After day 9, however, their ΔUI decreased by ∼50% and no longer differed from WT. In contrast, while ‘Ob-#6’ was comparable to ‘Ob-Low’ in the initial phase (2.5ΔUI/mouse/day), ΔUI increased 11.5-fold to WT between day 9-27 (8.3ΔUI/mouse/day), which was comparable to ΔUI of ‘Ctrl-Ob’ (12.3±1.0ΔUI/mouse/day). After day 28, the ΔUI of ‘Ob-#6’ further increased 23-fold over WT, however, this was co-caused by water spillage through the stereotypic grooming behavior. The water intake mirrored the UI results, with ‘Ob-Low’ exhibiting an initial increase (day 2: 5.8±0.2ml/mouse/day, peak day 14: 10.4±0.4ml/mouse/day), followed by a gradual decrease (day 44: 6.1±0.4ml/mouse/day, Fig. 2k). Consistently, ‘Ob-#6’ diverged from ‘Ob-Low’ and progressively increased between day 9-28 (day 9: 10.1ml/mouse/day), until stabilizing the water intake after day 28 (between 24-29ml/mouse/day). Correspondingly, the blood glucose revealed that ‘Ob-#6’ was the only mouse that developed severe hyperglycemia, and the polyuria onset coincided with the point at which blood glucose consistently exceeded 20mM (Fig. 2l), suggesting a threshold effect. While ‘Ob-Low’ occasionally had blood glucose levels approaching or exceeding 20mM, most measurements showed only mild to moderate hyperglycemia (range: 10.1±2.9mM ‘Ob-#1’ to 15.0±3.6mM ‘Ob-#9’, Fig. 2l-m).

### STZ-Pee pilot

To explore the generalizability of the UI, we employed data of a streptozotocin (STZ)-induced T1DM pilot with 10 female WT, 5 receiving STZ and 5 control (Ctrl) treatment (2&3 mice/cage/group). In week 4, the blood glucose response was strongest in both mice of cage ‘STZ-1’ (22.9±2.0mM), while the response in ‘STZ-2’ was only mild (15.8±1.3mM, Fig. 2n). Interestingly, the relatively small difference of mean blood glucose between the two STZ cages was associated with a striking difference in UI (Fig. 2o). Compared to Ctrl (1.3±0.2UI/mouse/day), UI was increased 2-fold in ‘STZ-2’ (2.5UI/mouse/day), but 9-fold in ‘STZ-1’ (11.6UI/mouse/day, Fig. 2p). Analysis of circadian urination revealed that polyuria in ‘STZ-1’ occurred throughout the 24-hour cycle, peaking in the middle of the dark phase, where feeding-induced hyperglycemia would be expected to amplify diuresis in severely hyperglycemic animals (Fig. 2q). Likewise, the UI of ‘STZ-2’ peaked in the late dark phase, but was virtually identical to WT during the rest of the circadian cycle, suggesting that these mice only become severely hyperglycemic postprandially. Again, the water intake mirrored the UI (Fig. 2r), with an increase of 1.9-fold in ‘STZ-2’ (7.6ml/mouse/day) and 7.8-fold in ‘STZ-1’ (31.8ml/mouse/day) compared to WT (4.1±0.3ml/mouse/day, Fig. 2s). Remarkably, food intake in ‘STZ-1’ was increased 2-fold (6.6kcal/mouse/day), while ‘STZ-2’ (3.5kcal/mouse/day) was virtually identical to WT (3.3±0.2kcal/mouse/day, Fig. 2t-u). Considering the relatively modest blood glucose difference between STZ cages, the disproportional potentiation of the three key symptoms of DM - polyuria, polydipsia, and polyphagia – appears extraordinary, and reinforce the notion of a threshold effect occurring when blood glucose exceeds 20mM.

### STZ-Pee 2.0

Encouraged by aforementioned observations, we performed a follow-up study with 12 Ctrl and 16 STZ-treated, pair-housed females for 8 weeks. Blood glucose (Fig. 3a), food (Fig. 3b) and water intake (Fig. 3c) were assessed twice weekly and UI was tracked continuously (Fig. 3d). Prior to treatment, UI was low and similar, thereafter it increased 12.6-fold in STZ (6.6±1.0UI/mouse/day vs. 0.5±0.05UI/mouse/day, p<0.0001, Fig. 3e). By inspection of the combined data for each cage, a dose-dependent effect of hyperglycemia on the other parameters was immediately apparent. In Ctrl cages, where blood glucose remained stable, UI, food, and water intake were also constant (Fig. 3f, Supplementary Fig. 1a-f). Conversely, the magnitude of the changes in food, water, and UI essentially reflected the average magnitude of the STZ-mediated hyperglycemia of both mice in a STZ cage (Fig. 3g-I, Supplementary Fig. 1g-n). Importantly, polyuria was clearly detectable, even if only one mouse in a cage developed hyperglycemia, indicating that the UI is also reliable for detection of individual cases of polyuria in group-housed mice (Fig. 3h). Given the variable response to STZ, a wide range of values was observed for all measured parameters (Fig. 3j-m). Consequently, we investigated the diagnostic potential of UI as a quantitative indicator of hyperglycemia, polydipsia, and polyphagia.

**Fig. 3:**
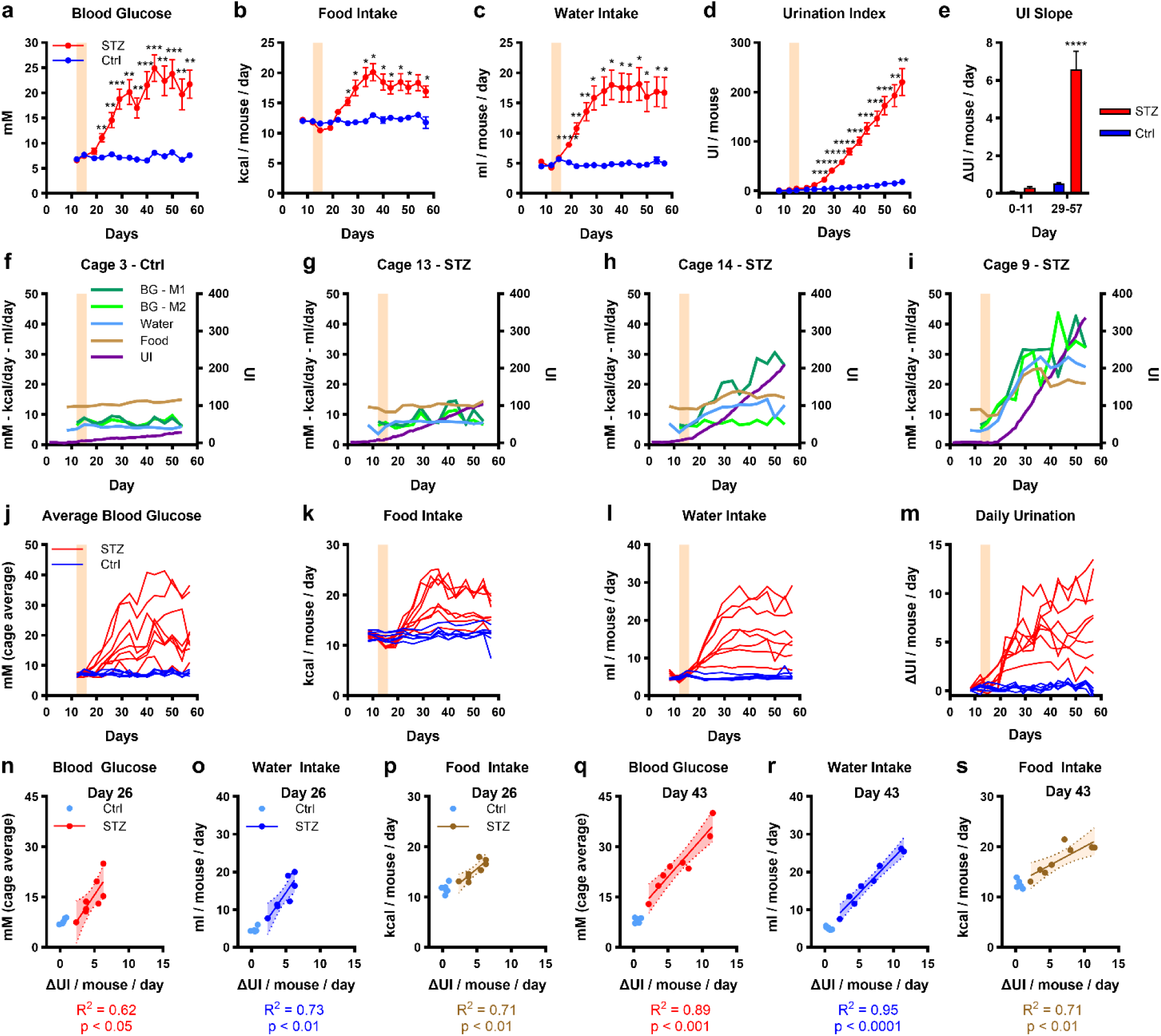
STZ-Pee 2.0 study - main effects and correlations. **a-d,** Time course development (mean±SEM) of blood glucose (**a**), food intake (**b**), water intake (**c**), and UI (**d**) following STZ dosing (shaded area) in control cages (n=6) and STZ-treated cages (n=8) of female C57BL/6 mice. **e**, Average daily UI before (day 0-11) and after STZ (day 29-57) (mean±SEM of UI slopes). **f-i**, Individual time course data from representative cages of control (M1: mouse-1, M2 mouse-2, **f**), low-responder STZ (**g**), single-responder STZ (**h**), and dual high-responder STZ (**i**). Blood glucose are from individual mice, while food intake, water intake, and UI are normalized for housing density. **j-m**, Time course data for individual cages of control (blue) and STZ (red) of blood glucose (cage average) **(j**), daily food intake (**k**), daily water intake (**l**), and daily UI change (**m**). **n-p**, correlations and 95% CI (shaded area) between daily UI change and blood glucose (cage average) (**n**), water intake (**o**), and food intake (**p**) in the STZ group 10 days after treatment. **q-s**, correlations and 95% CI (shaded area) between daily UI change and blood glucose (cage average) (**q**), water intake (**r**), and food intake (**s**) in the STZ group 27 days after treatment. *p<0.05; **p<0.01; ***p<0.001; ****p<0.0001.

Blood glucose positively correlated with UI already on day 22, 6 days post-STZ (Supplementary Fig. 2a). On day 26, we found strong positive correlations between UI and blood glucose, water, and food intake (Fig. 3n-p), which even increased by day 43 (Fig. 3q-s). In fact, the linear relationship between UI and blood glucose, water, and food intake remained strong from day 26 until end, except for day 33 (Supplementary Fig. 2a-c). The polyphagia, observed in ‘STZ-#1’ in the pilot, was recapitulated in 50% of STZ cages (Supplementary Fig. 3a). These also showed the highest increase of blood glucose (Supplementary Fig. 3b), UI (Supplementary Fig. 3c), and water intake (Supplementary Fig. 3d). Only food intake exhibited this seemingly dichotomous response pattern.

### Subdivision of the STZ-2.0 Responders

Considering the differing response to STZ, we split the STZ group into low-responder and high-responder. While caloric intake increased by ∼20% in ‘STZ-Low’, polyphagia was indeed severe in ‘STZ-High’, peaking about twice of ‘Ctrl’, with both STZ groups becoming significant from day 26 (Fig. 4a). Again, the main driver appears to be whether blood glucose exceeds 20mM, as this was a consistent differentiator between groups after day 26 (Fig. 4b). This supports a tipping point existence at 20mM. Although similar trajectories were found for glucose (p<0.001, Fig. 4b), daily water intake (p<0.0001, Fig. 4c) and urination (p<0.0001, Fig. 4d), temporal dynamics differed. Glucose was elevated from day 22 in ‘STZ-High’ (Fig. 4, red arrow) and from day 29 in ‘STZ-Low’ (Fig. 4, blue arrow). In contrast, polydipsia emerged significantly in both groups from day 19, only 3 days post-STZ (Fig. 4c). Summarized according to the temporal resolution of the other metrics, daily UI significantly increased on day 22 (Fig. 4d). However, the UI in 1-day resolution revealed a significant increase in both STZ groups already on day 20 (Fig. 4e). From day 29-57, the average UI increased 8.6-fold in STZ-Low (4.5±0.6UI/mouse/day) and 16.6-fold (8.7±0.8UI/mouse/day) in ‘STZ-High’ compared to ‘Ctrl’ (Fig. 4f). Thus, water intake and UI were the most sensitive indicators of developing hyperglycemia, with UI outperforming manual glucose measurements in STZ groups by 2 and 9 days, respectively. In fact, with the longest latency from treatment to measurable effect in STZ-low, intermittent glucose testing turned out to be our least reliable metric. This is no coincidence considering that glucose measurements only reflect the glycemic status exactly at sampling, i.e. usually during light phase. Comparing circadian urination patterns revealed the limitation of studying hyperglycemic models only in light phase (Fig. 4g-i). Before and during STZ treatment, UI patterns were similar (Fig. 4g-h). However, the circadian profile post-STZ revealed a biphasic pattern, with most of the increased UI occurring in the first 9 hours of dark phase, followed by a second at beginning of light phase (Fig. 4i). Investigating the weekly progression of circadian UI, we found that dark-phase polyuria developed in both groups one week after treatment (Supplementary Fig. 3e). In the second week, the separation between ‘STZ-High’ and ‘STZ-Low’ emerged at beginning of dark phase (Supplementary Fig. 3f). A full separation, including early light-phase polyuria, was established by week 3 post-STZ and remained the subsequent 3 weeks (Supplementary Fig. 3g-j).

**Fig. 4:**
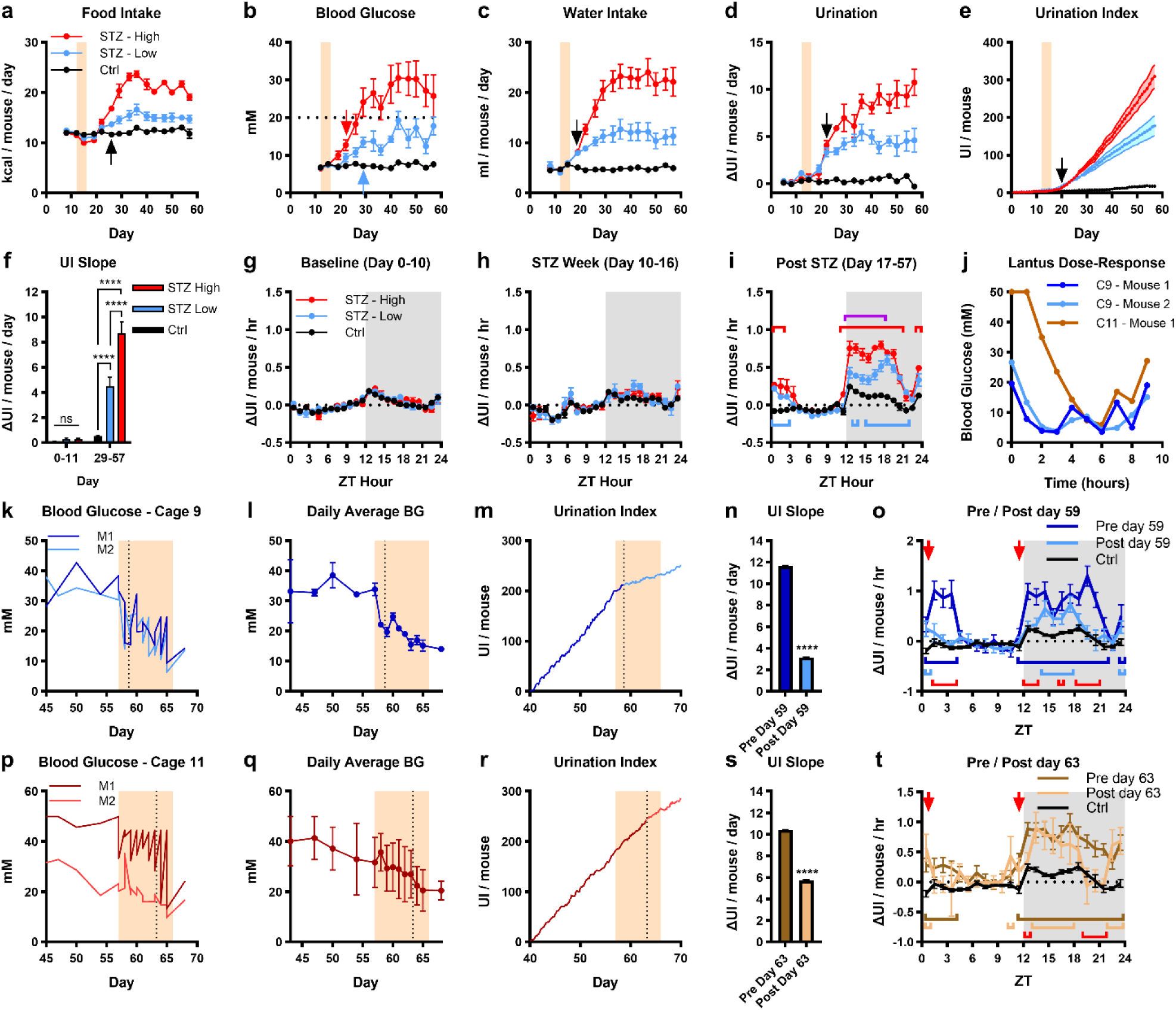
STZ-Pee 2.0 - High/low responder analysis and pharmacological intervention. **a-e,** Time course development (mean±SEM) in STZ-high responder (n=4) and STZ-low responder (n=4), and control (Ctrl) cages (n=6) of food intake (**a**), blood glucose (**b**), water intake (**c**), daily UI change (**d**) and UI (**e**) following STZ dosing (shaded area). Main effects and interaction between group and time were highly significant for all of **a-e**. Arrows denote the first significant divergence of STZ groups from control (p < 0.05). Black arrows indicate simultaneous divergence of STZ-High and STZ-Low from Ctrl, while divergence at different time points is indicated in arrows matching the group color. All other indicators of significance have been omitted for clarity. **f**, Average daily urination rate before (day 0-11) and after STZ (day 29-57) (mean±SEM). **g-i**, Circadian change in UI (mean±SEM) before STZ (**g**), during STZ treatment (**h**), and after STZ (**i**). Significant differences from control for each STZ group are indicated with brackets in the group color, and purple bracket indicates differences between STZ-High and STZ-Low. **j**, Blood glucose during insulin Lantus dose-response test (1U/mouse). **k-t**, Lantus intervention in cage-#9 (blue) and cage-#11 (brown). Time course of individual blood glucose (**k, p**), daily average blood glucose (mean±SEM) (**l, q**), and UI (**m, r**). Shaded area is the treatment period, and dashed line indicates time point of UI slope change. **n**, **s**, Average daily urination rate before and after UI slope change (slope±SE). **o**, **t**, Circadian change in UI before and after UI slope change compared to control cages (n=6). Red arrows show time of Lantus dosing, brackets in group colors indicate significant differences from control, and red brackets indicate differences between pre and post. *p<0.05; **p<0.01; ***p<0.001; ****p<0.0001.

### UI reversal by pharmacological hyperglycemia treatment

Having established the reliability of the UI for monitoring onset and development of polyuria, we evaluated its applicability to assess the therapeutic efficacy of glucose-lowering intervention with long-acting insulin, for mice with constant glucose >30mM in week 4-6 (both ‘cage-9’, one ‘cage-11’). An initial dose-response test to validate effective glucose lowering exhibited that both ‘cage-9’ mice remained around the euglycemic range (BG <10mM) for 8 hours after dosing, while the treated ‘cage-11’ mouse took 4 hours to reach comparable levels due to the initial >50mM blood glucose (Fig. 4j). During the subsequent treatment period, glucose, measured at dosing time, was effectively reduced in both ‘cage-9’ mice (Fig. 4k), by over 50% (day 57: 33.8±1.4mM, day 65: 15.2±1.3mM, Fig. 4l). After two days of treatment, the UI trajectory changed substantially (Fig. 4m), corresponding to a 73% reduction (Fig. 4n), and coinciding with the point where glucose reached ∼20mM. This was driven by an overall decrease in the circadian urination pattern, with dark phase polyuria reduced from 11 to 5 hours and light phase polyuria essentially eliminated (Fig. 4o). In ‘cage-11’, treatment was also effective but showed higher variability and a distinct pattern of lower afternoon glucose compared to morning (Fig. 4p). During treatment, glucose of both mice gradually declined by 35% (day 57: 31.7±7.0mM, day 65: 20.5±5.9mM, Fig. 4q). The UI trajectory changed when average glucose reached ∼25mM (Fig. 4r), corresponding to a 45% UI reduction (Fig. 4s), which is remarkable given that only one mouse was treated. Again, light phase urination normalized almost completely to ‘Ctrl’, while dark phase polyuria was blunted and reduced from 12 to 7 hours (Fig. 4t).

## Discussion

Our study demonstrates accurate detection of hyperglycemia by continuous bedding monitoring in an automated home cage system, representing a significant advance in metabolic research methodology. This approach addresses critical challenges of traditional glucose measurements, which are invasive, time-consuming and only intermittent snapshots of glucose levels^10^. The strong correlation between bedding moisture and blood glucose during hyperglycemia validates our metric as reliable biomarker, particularly in severely diabetic mice, showing up to 8.5-fold increased UI.

The correlation between UI and other metabolic parameters reflects the DM pathophysiology, in which polyuria is accompanied by hyperglycemia, polydipsia and polyphagia^16^. Our multi-parameter monitoring approach provides insights into the complex diurnal rhythm of hyperglycemia and its effects on voiding behavior^17,18^. To capture these patterns, especially during dark phase, is important given that voiding behavior regulation is known to be altered in DM through various mechanisms, including vasopressin-induced aquaporine-2 upregulation^35^ and prostaglandin E2-mediated effects^36^. This is particularly useful for monitoring (large) colonies that develop spontaneous hyperglycemia, such as the non-obese diabetic mice, which traditionally require frequent sampling between 12-30 weeks of age^37^. This aligns with current efforts to improve continuous glucose monitoring in animal models while reducing sampling stress^4,7^.

The non-invasive UI significantly enhances animal welfare by eliminating stress, and stress-induced blood glucose fluctuations, associated with traditional sampling methods^5,38^. Handling-free sampling can be done via catheterization^39^ or CGM, but requires surgical expertise, specialized setup and skilled personnel, which either is still limited to glucose snapshots or, for CGM, needs calibration with handling and can affect glucose regulation^11,12^. Our approach circumvents such limitations while providing continuous data. Our findings demonstrate the method’s sensitivity to therapeutic interventions, as evidenced by successfully reversing the polyuria phenotype following pharmacological insulin treatment of hyperglycemic STZ mice. Tracking treatment responses in real-time enables precise monitoring of intervention efficacy or timing of treatments according to the onset of hyperglycemia, potentially reducing model variability and improving research outcomes.

A key advantage of our approach is its group-housing compatibility, avoiding alterations associated with individual housing, e.g. required in metabolic cage phenotyping. Individual housing has been shown to affect energy intake and expenditure, potentially inducing anxiety- and depression-like behavior that could confound experimental results^14,40^. Monitoring polyuria in group-housed mice in their home cage minimizes adverse effects of human presence and environmental changes^41^. Moreover, the DVC simultaneously collects data beyond BSI, such as circadian activity or changes of sleep patterns^32,42^, enabling multi-faceted characterization with more complex data compared to blood glucose assessment alone. This might highlight novel aspects or effects of treatments or genotypes. Integrating this methodology with existing DVC technology builds upon established applications in animal husbandry management and welfare monitoring^43,44^. This approach extends beyond traditional void spot assays and metabolic cage analyses, which induce artifacts and stress in laboratory animals^23,24^.

However, several limitations warrant consideration. UI’s susceptibility to artifacts from water spillage or bottle leakage during cage-handling necessitates careful monitoring and documentation of maintenance tasks. Furthermore, as evaporation characteristics vary with bedding materials and amounts, controlled tests should be performed to identify the optimal bedding amount to minimize evaporation variations. Particularly in models with only mildly increased urination, evaporation can be a serious confounder, especially if bedding is not held constant. Another important consideration for UI tracking when sharing a DVC rack for multiple experiments are cage and position changes as external, unrecognized activities can affect the own cages. Therefore, all (cage) parameters should be accurately captured and kept consistent during UI recordings, which in turn benefits the data quality. Additionally, while the correlation between blood glucose and UI is robust, the detection sensitivity may be reduced in group-housed animals with only mild hyperglycemia, similar to challenges facing with other glucose measuring systems^15^.

In conclusion, our innovative approach enables a sample-free assessment of hyperglycemia by deriving the digital biomarker UI from BSI. This provides a non-invasive, continuous method for tracking disease onset and progression as well as treatment efficacy. Detecting responses and effects of treatments and other (therapeutic) interventions renders UI highly relevant, especially for pharmacological studies. Maintaining social housing combined with simultaneously assessment of multiple parameters, makes it a useful tool for large-scale studies and longitudinal monitoring of disease progression. Our methodology represents a significant advance regarding the 3R principles for diabetes research, improving animal welfare and data quality.

## Methods

### Ethical statement

All experiments were performed in accordance with Guidance on the Operation of the Animals (Scientific Procedures) Act 1986 and associated guidelines, EU Directive 2010/63, complied with institutional ethical and ARRIVE guidelines and have been authorized by the responsible national authorities. STZ experiments were approved by Landesamt für Gesundheit und Soziales Berlin (G0104/20) and performed at the Forschungseinrichtung für Experimentelle Medizin (FEM) at Charité - Universitätsmedizin Berlin. The INSPIRE cohort was carried out in accordance with the French Ministry of Agriculture and Toulouse University ethic committee and approved by the Ministry of Superior Education and Research (APAFIS2019120614331282, APAFIS2020022409196014). Ob/ob mice for urination observations were ordered on license (2019-15-0201-00073).

### DVC monitoring and husbandry conditions

Mice were predominantly group-housed (acclimatization: 7-14 days), fed ad libitum and maintained under standardized site-specific husbandry conditions with 22±2⁰C, 55±10% RH, a 12-hour light/dark cycle (06:00/18:00) in DVC racks (Tecniplast) for data recording. Cohort-specificities are indicated for CD1, ob/ob (SAFE D30 chow, SAFE Aspen bedding), INSPIRE^34^ and STZ^32^ (SAFE FS 14 bedding). As enrichment, cages were supplied with transparent tunnels, translucent red shelters, bite sticks, nesting material and standardized bedding amounts. The DVC home cage-based monitoring system non-invasively tracks electrode data of each cage in a DVC rack via capacitance sensing technology (CST) 24/7 in real time. A sensor board containing 12 electrodes under each cage records changes in the electromagnetic fields once every 0.25s^26^. For our study, we export the BSI data and processing to acquire the Urination Index (UI).

### “In vitro” water and diet tests and procedures

In vitro tests and procedures are detailed in Supplementary Methods.

### Animal models

#### INSPIRE cohort

Voiding behavior data of the INSPIRE cohort was assessed at ages 6, 12, 18, and 24 months^33,34^. We evaluated the DVC data of 123 male and 87 female cages for UI 30 days prior to voiding tests, and compared the 30-day UI change to the total area of urine spots.

#### Housing density analysis

As a preliminary assessment of UI linearity, we retrospectively analyzed historical data from a 30-week-old CD1 stock colony (Crl:CD1(ICR), Charles River) housed in the DVC for data recording. Housing densities were 1, 2, 3, 4, or 5 males or females per sex and cage. During the recording (34 days), only routine husbandry procedures were performed.

#### Ob/Oh-Pee study

As T2DM model, we analyzed DVC data from 2 individually housed, 25-week-old, male ob/ob mice (B6.V-Lep^ob^/JRj, Janvier Labs), which were part of an unrelated study. We included their DVC and blood glucose data that was consistently above the reading range of handheld glucometers (>33.3mM, Contour XT/Next, Bayer), providing a model of severe uncontrolled T2DM. During the recording (23 days), only routine husbandry procedures were performed.

To investigate the time course of phenotype development analogous to these ob/ob mice, we performed a longitudinal study as a urination reference with 10 male ob/ob and 16 male and 16 female wildtype (WT) mice starting at 3 weeks of age. Ob/ob males were individually housed in the DVC to enable quantitative comparison of blood glucose and UI. Glucose from tail vein pricking, body weight, and water consumption by manual bottle weighing were recorded 3-times per week at 10 am. Starting around day 27, one ob/ob mouse (Ob-#6) developed a stereotypic behavior of excessive grooming on the nozzle of the bottle, causing water to spill in the cage. The WT reference mice (C57BL/6JRj, Janvier Labs) were group-housed as 4 per cage, only recording DVC data without assessing glucose, body weight, or water consumption.

#### STZ-Pee pilot and STZ-Pee 2.0

To induce polyuria, we used the STZ-mediated T1DM model with C57Bl/6J females (Charles River) for both studies. In the STZ-Pee pilot, 10 8-week-old females were housed in 2 cages with 2 mice and 2 cages with and 3 mice in the DVC. At 10 weeks, 5 mice (2 cages) were injected with vehicle-control (Ctrl, buffer: 0.1M citric acid: 0.1M Na-citrate 3:2, pH=4.0) and 5 with STZ (55mg/kg, freshly dissolved and used within 15min) on 5 consecutive days for pancreatic islet β-cell destruction. Due to expected polyuria, cages were regularly changed twice a week with monitoring of diet and water intake. Blood glucose was monitored in week 0, 4 and 8, followed by sacrificing for organ harvest.

For STZ-Pee 2.0, 28 5-week-old females were pair-housed in the DVC system, and at 7 weeks of age, 12 mice (6 cages) were injected with Ctrl and 16 mice (8 cages) with STZ for 5 consecutive days. Cage changes were standardized twice a week, on Monday and Thursday. At same days, mice were scored and monitored for body weight, diet and water intake and blood glucose measurements in the morning. After reaching constant hyperglycemia (>30mM) in week 4-6, respective mice were treated by intraperitoneal injection of insulin glargine (1U, Lantus, Sanofi, Germany) twice daily between 6am and 7am and before start of night phase between 5pm and 6pm. Both mice from cage-#9 and one of cage-#11 met these inclusion criteria. The other mouse in cage-#11 did not and was therefore not treated. Finally, mice were sacrificed for organ harvest.

### Statistical and data analyses

Data processing, graph generation and statistical analyses were performed and generated in Excel (Microsoft), Prism 9 (GraphPad) and the Urinator App. All data are presented as the mean±SEM unless indicated otherwise for cage data or individual data points for correlations.

Data was analyzed as one-way ANOVA, group-wise comparison by ANOVA, repeated-measures two-way ANOVA adjusted with Bonferroni multiple comparisons test for post hoc analysis or correlations and 95% CI as indicated. Sample size or cage number are included accordingly in each figure legend. Statistical significance was set as *P≤0.05, **P≤0.01, ***P≤0.001, ****P≤0.0001.

### App design and workflow

The Urinator app was coded in R (v4.3.1 & shiny v1.9.1), available on https://github.com/Mortendall/UrinatoR, and hosted on https://cbmr-rmpp.shinyapps.io/UrinatoR/. Package dependencies and version control were managed with renv (v1.0.7).

App workflow is as follows: After assigning a CSV separator and local decimal mark, the app pre-processes an uploaded CSV file by standardizing column names and removing summary statistics. Time stamps are converted to local time and groups are inferred based on column name. Next, an event list (csv) only filtering cage INSERTION events is uploaded. Each event is assigned to the closest matching data point using the fuzzyjoin package (v0.1.6). Delta value for each time point X is calculated as value(x)=rawdata(X)−rawdata(X+1), and delta values are excluded in a pre-determined window around each event to account for water bottle spillage from cage handling, and to exclude cage changes. Once delta values have been calculated and data has been trimmed, the user inputs the number of mice per cage to calculate increases per mouse. Cumulative values are calculated for each cage/mouse. Summary stats are generated, and DVC signals can be visualized via Plotly (v4.10.4) as raw data, cumulative values or incremental changes for individual cages or experimental groups. Urination can also be visualized per hour for experimental groups or individual cages. All data can be downloaded as an xlsx file.

## Reporting summary

Further information on research design is available in the Nature Portfolio Reporting Summary linked to this article.

## Data availability

All data analyzed or generated in this study are included in the main text or the supplement. The datasets are available from the corresponding author upon reasonable request.

## Acknowledgements

We thank Diana Woellner, Marie-Christin Gaerz and Nadine Huckauf, Charité – Universitätsmedizin Berlin for excellent assistance with STZ experiments and Bianka Verret and her animal caretaker team (FEM) at the MRC, Charité – Universitätsmedizin Berlin. We also thank Kristoffer Egerod and Nadia Nielsen Aalling for offering ob/ob mice for home cage monitoring, and Christine Broholm and Carina Onuczak Rosenberg for their technical expertise and assistance throughout the mouse studies.

## Author information

These authors contributed equally: Sebastian Brachs, Morten Dall.

These authors jointly supervised this work: Stefano Gaburro, Thomas Svava Nielsen.

## Contributions

S.B., T.S.N. and S.G. conceived the project. S.B., M.D. and T.S.N. designed the study. S.B., M.D., L.-K.Z., Y.S., C.O., A.P. and T.S.N. planned, conducted, supervised animal experiments, collected and/or provided data. M.D. and T.S.N conducted data analysis. K.M. provided funding, facility resources and laboratory space. T.S.N. performed statistical testing and constructed figures. S.B., M.D., L-K.Z., S.G. and TS. N. wrote the manuscript with input from all authors.

## Ethics declarations

