## Supplemental methods and figures for "Robust non-invasive detection of hyperglycemia in mouse models of metabolic dysregulation: the novel Urination Index biomarker"

### Supplementary Methods

#### "In vitro" water and diet tests

To evaluate the relationship between UI and bedding moisture, a dose-response test was performed by injecting 1, 2, 5, and 10ml water in cages containing bedding but no mice. Likewise, we tested signal sensitivity to common diets, chow or 60% high-fat diet (HFD, D12492i, Research Diets), by adding 1, 2, and 4 pellets to the cages. To assess whether the amount of aspen bedding impacted the signal, we performed all tests twice in parallel in cages with different bedding amounts.

For water tests, two groups of 6 cages each were prepared with either 100g or 200g standard bedding (SAFE aspen) and inserted in the DVC system. Each bedding group was split into a small-bolus and large-bolus group (each n=3). After 30min of baseline recording, a water bolus of 1ml was injected in the small-bolus cages, 5ml large-bolus cages through the water bottle opening. After 15min of data collection, additional 2ml bolus and 10ml were injected for small-bolus and large-bolus cages. Following this, data was recorded for 3 days to evaluate the rate of change in UI during the evaporation of 3ml and 15ml of water, respectively.

On day 4, diet pellets were added to the cage floor to evaluate the contribution of spillage or shredding to changes in UI. Chow pellets were added to the small-bolus and HFD pellets to the large-bolus cages. In all cages, pellets were added three times separated by 30min of recording, starting with 1, then 2, and finally 4 pellets. Due to density differences between chow and HFD, the mass of pellets added in each step was different. Consequently, all analyses were done according to the measured pellet mass to establish the effect per gram of diet.

**"In vitro" procedure tests**

Since water bottles for rodents generally tend to drip when handled, we decided to test the extent of water spillage caused by cage handling. Therefore, we set up 2 cage groups (each n=8) equipped with bottles, nesting material, shelter, and either 100 or 200g bedding. Water bottles were filled ~50% to model a worst-case scenario, since the risk of leakage increases if bottles are less than full. Following a 30-min baseline recording, we simulated a typical cage-handling event, where each cage was removed from the rack, placed on a trolley, the bottle was removed, placed nozzle-up, the cage was opened and closed. Then, the bottle was re-inserted in the lid and the cage was returned to the same DVC slot. After return of the last cage, data was recorded for 15 min to acquire post-handling signals.

Finally, we tested the effect of cage insertion/removal above and below the experimental cages. Since the DVC electrodes are omnidirectional, we speculated that cage insertion/removal below, where the bottle is located immediately underneath the sensor board of the cage above, could be detected as a UI change in the upper cage. Conversely, cage insertion above should not affect the UI. Hence, after a 15-min baseline recording, we inserted cages with full bottles either above or below the test cages, recorded for 15min, and removed them, followed by another 15-min recording.

### Supplementary figures

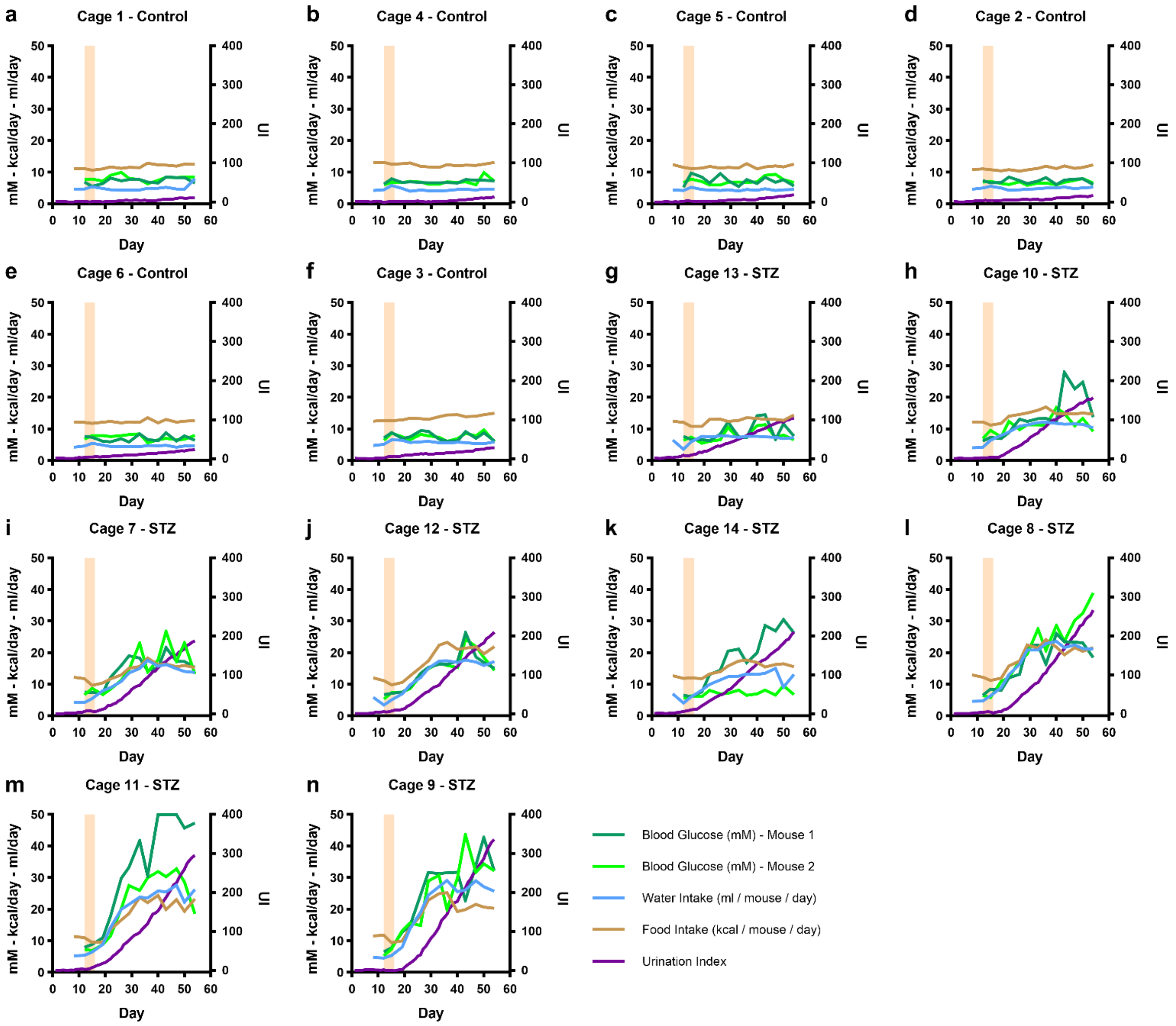


**Supplementary Fig. 1: STZ-Pee 2.0 - Individual time course data from all cages.** Blood glucose from both mice and food intake, water intake, and UI, which were all normalized for housing density, in control cages (**a-f**) and in STZ treated cages (**g-n**). STZ treatment is indicated by the shaded area. Blood glucose (mM), food intake (kcal/mouse/day) and water intake (ml/mouse/day) are on the same scale and thus plotted together on the left axis, but with different units. UI is plotted on the right axis.

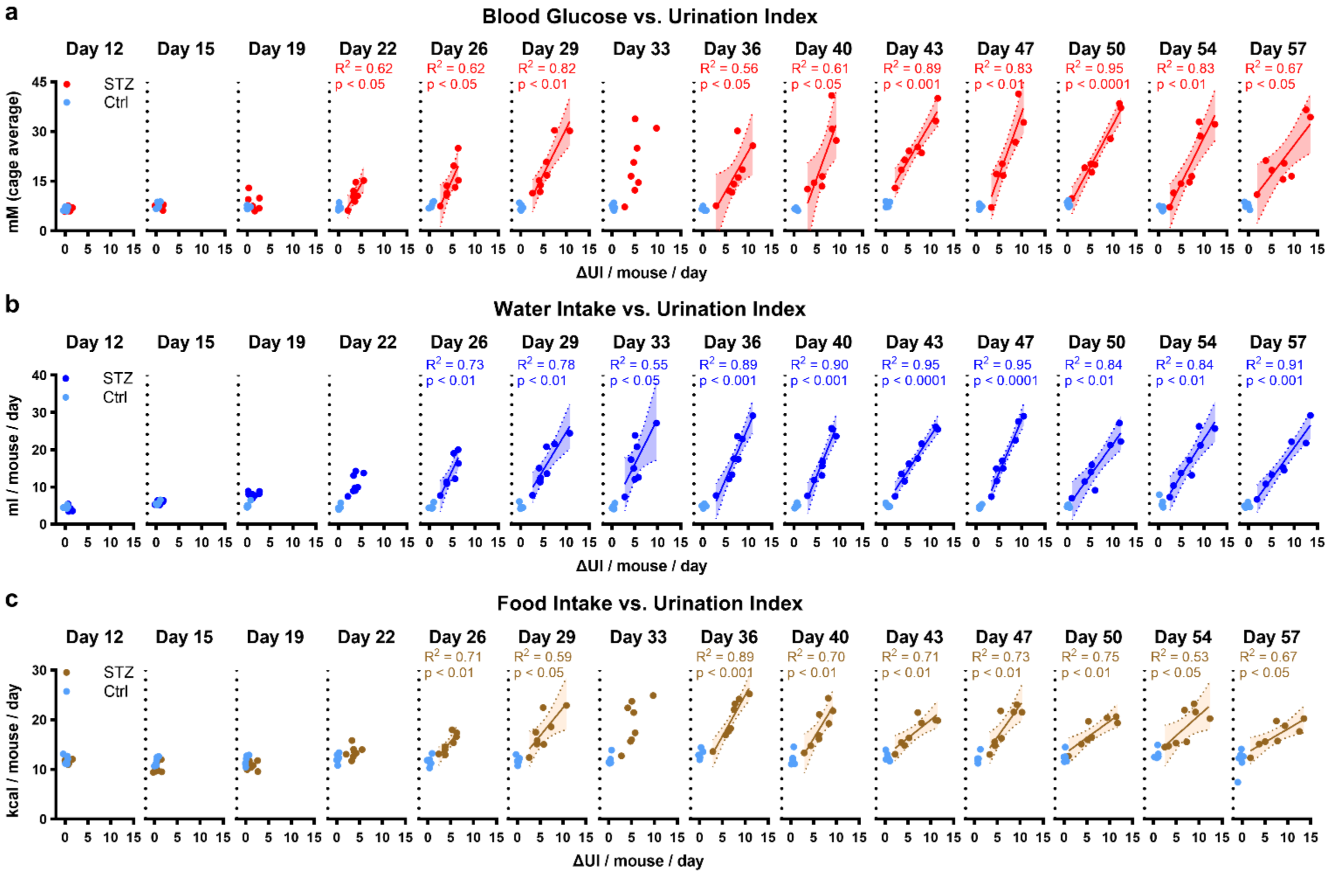


**Supplementary Fig. 2: Correlation time course for UI with blood glucose, water intake and food intake.** Correlation analysis was performed for STZ-treated cages at all time points where blood glucose, water intake and food intake were determined. 95% CI (shaded area), $R^2$, and p-value are included for significant correlations only ($p<0.05$).

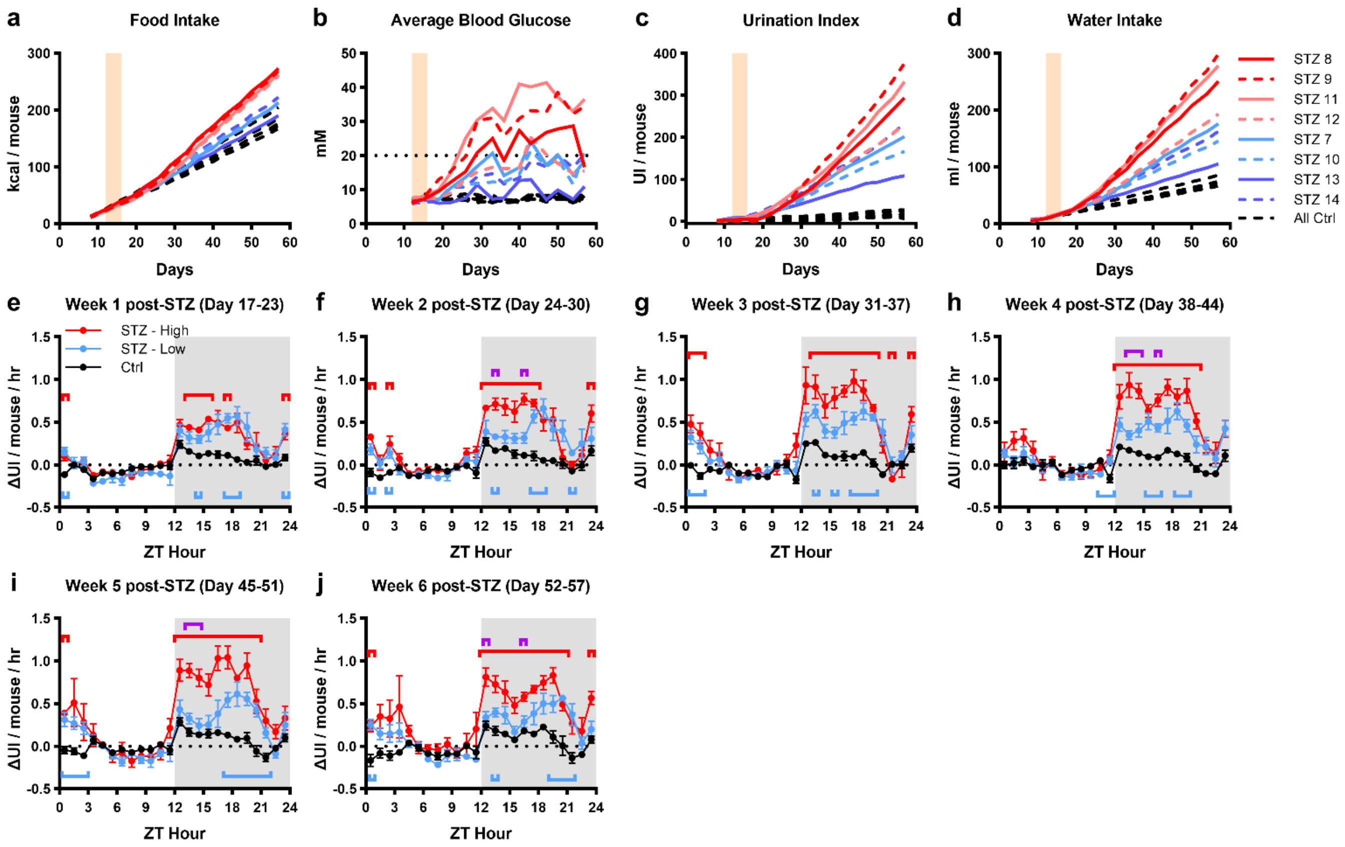


**Supplementary Fig. 3: STZ-Pee 2.0 - High/Low Response Analysis. a-d**, time course of cumulative food intake (**a**), average blood glucose (**b**), UI (**c**), and cumulative water intake (**d**) for individual cages grouped according to whether polyphagia was apparent (red, STZ high-responder) or not (blue, STZ low-responder) relative to control (black). Each STZ cage is shown with unique line color/style to highlight the consistency in responses across different metrics. **e-j**, Weekly circadian profiles of UI changes (mean±SEM) following STZ treatment in STZ-high responder (n=4) and STZ-low responder (n=4), and control cages (Ctrl, n=6). Significant differences from control for each STZ group are indicated with brackets in the group color, and purple bracket indicates differences between STZ-High and STZ-Low ($p<0.05$).
